# Different neural networks for conceptual retrieval in sighted and blind

**DOI:** 10.1101/384552

**Authors:** Roberto Bottini, Stefania Ferraro, Anna Nigri, Valeria Cuccarini, Maria Grazia Bruzzone, Olivier Collignon

## Abstract

We investigated the experiential bases of knowledge by asking whether people that perceive the world in a different way also show a different neurobiology of concepts. We characterized the brain activity of early-blind and sighted individuals during a conceptual retrieval task in which participants rated the perceptual similarity between color and action concepts evoked by spoken words. Adaptation analysis showed that word-pairs referring to perceptually similar colors (e.g., red-orange) or actions (e.g., run-jump) led to repetition-suppression in occipital visual regions in the sighted, regions that are known to encode visual features of objects and events, independently of their category. Early blind showed instead adaptation for similar concepts in language-related regions, but not in occipital cortices. Further analysis contrasting the two categories (color and action), independently of item similarity, activated category-sensitive regions in the pMTG (for actions) and the precuneus (for color) in both sighted and blind. These two regions, however, showed a different connectivity profile as a function of visual deprivation, increasing task-dependent connectivity with reorganized occipital regions in the early blind. Overall, our results show that visual deprivation changes the neural bases of conceptual retrieval, which is partially grounded in sensorimotor experience.

**Significance Statement:** Do people with different sensory experience conceive the world differently? We tested whether conceptual knowledge builds on sensory experience by looking at the neurobiology of concepts in early blind individuals. Participants in fMRI heard pairs of words referring to colors (e.g., green-blue) or actions (e.g., jump-run) and rated their perceptual similarity. Perceptual similarity of colors and actions was represented in occipital visual regions in the sighted, but in language-related regions in the blind. Occipital regions in the blind, albeit not encoding perceptual similarity, were however recruited during conceptual retrieval, working in concert with classic semantic hubs such as the Precuneus and the lpMTG. Overall, visual deprivation changes the neural bases of conceptual processing, which is partially grounded in sensorimotor experience.

## Introduction

As we think, we navigate and retrieve conceptual knowledge, but the nature of this knowledge is highly debated. One of the major sources of disagreement is whether thinking is more similar to the analog replay of our experience (Barsalou, 1999; Kosslyn, Ganis, & Thompson, 2001), or the symbolic computations of a Turing machine (Fodor, 1975; Pylyshyn, 1984).

Congenital blindness offers an ideal model to test whether thinking is simulating our own experience of the world (Barsalou, 2009; Lakoff & Johnson, 1999). If conceptual processing is largely grounded into experience, blind people, who experience the world in a different way, should also show a different neurobiology of concepts, at least in part (Casasanto, 2011). Several studies, however, seem to provide evidence against this idea (Bedny & Saxe, 2012). For instance, when sighted and blind were asked to retrieve information about highly visual entities, knowledge about small and manipulable objects (Peelen et al., 2013) activated the lateral temporal-occipital complex; thinking about big non-manipulable objects activated the parahipoccampal place area (He et al., 2013); and processing action verbs (compared to nouns) activated the left posterior middle temporal gyrus (Bedny et al. 2012) in both groups. These results seem to suggest that blindness leaves the neurobiology of conceptual retrieval largely unchanged, and that experience plays a minor role in shaping mental representations (Bedny & Saxe, 2012; Uta Noppeney et al., 2003).

Alternatively, it is possible that previous studies relied on paradigms designed to investigate categorical knowledge, a level of processing that is relatively resilient to visual deprivation (Van Baelen, and Op de Beeck 2017). Indeed, categorical boundaries (e.g., between objects, tools or animals) are drawn based on several dimensions that exceed visual appearance (e.g., tokens from the same category do not always look alike; Bracci and Op de Beeck 2016; Proklova, Kaiser, and Peelen 2016), and depend on functional, emotional and linguistic constraints (Peelen & Downing, 2017). Indeed, brain regions that show category specificity, seem to represent different categories in a format that is at least partially independent from visual appearance (Bracci & Op de Beeck, 2016), and thus more likely to be resilient to visual deprivation (Bedny et al. 2012; van den Hurk, Van Baelen, and Op de Beeck 2017; Wang et al. 2015).

Differences between sighted and blind may be more likely to emerge when participants process the perceptual similarity of retrieved concepts (Martin, Douglas, Newsome, Man, & Barense, 2018), a process known to be encoded in more posterior (visual) occipital regions in the sighted (Fernandino et al. 2015; Borghesani et al. 2016; Mitchell et al. 2008). To test this hypothesis we asked sighted and early blind participants to rate pairs of concepts based on their perceptual similarity and analyzed the data using a repetition suppression framework (Barron, Garvert, and Behrens 2016; Wheatley et al. 2005; Grill-Spector, Henson, and Martin 2006). We reasoned that this method will allow to tag selectively the representation of low-level perceptual features during conceptual retrieval (Horner & Henson, 2011; Mohr, Linder, Linden, Kaiser, & Sireteanu, 2009). We predicted that perceptually similar concepts (within a given category) might elicit neural adaptation in the occipital cortex of sighted (who will retrieve their visual similarity), but not blind people. On the other hand, a direct comparison between categories (independently of visual/perceptual similarity) should highlight category-specific responses in more anterior/multimodal areas of the brain, with limited differences between sighted and blind. This should be especially true for conceptual categories that can be perceptually experienced both by sighted and blind individuals. Therefore, in our experiment, we choose stimuli exemplars coming from actions and color categories. Including colors, that can be experienced through vision only, allowed us to test whether their different epistemological status (concrete vs abstract), in sighted and blind, would influence their representation even in multimodal areas of the brain that usually show resilience to visual deprivation (Striem-Amit, Wang, Bi, & Caramazza, 2018).

## Materials and methods

### Participants

Thirty-six participants took part to this experiment: 18 early blinds (EB; 8F) and 18 sighted controls (SC; 8F). Participants were matched pairwise for gender, age, and years of education (Table S1).

All the blind participants lost sight at birth or before 3 years of age and all of them reported not having visual memories (Table S2). All participants were blindfolded during the task. The ethical committee of the Besta Neurological Institute approved this study (protocol fMRI_BP_001) and participants gave their informed consent before participation.

### Stimuli

We selected six Italian color words (rosso/*red*, giallo/*yellow*, arancio/*orange*, verde/*green*, azzurro/*blue*, viola/*purple*), and six Italian action words (pugno/*punch*, graffio/*scratch*, schiaffo/*slap*, calcio/*kick*, salto/*jump*, corsa/*run*). Words were all highly familiar nouns and were matched across categories (color, action), by number of letters (Color: mean= 5.83, sd= 0.98); Action: mean= 6, sd= 1.23), frequency (Zipf scale; Color: mean= 4.02, sd= 0.61; Action: mean= 4.18, sd= 0.4), and orthographic neighbors (Coltheart’s N; Color: mean= 14, sd= 9.12; Action: mean= 15.33, sd= 12.42).

Auditory files were made using a voice synthesizer (talk to me), with a female voice, and edited into separated audio files with the same auditory properties (44100 Hz, 32 bit, mono, 78 dB of intensity). The original duration of each audio file (range 356 – 464 ms) was extended or compressed to 400 ms using the PSOLA (Pitch Synchronous Overlap and Add) algorithm and the sound-editing software Praat (Boersma & Weenink, 2018). All the resulting audio files were highly intelligible.

### Experimental Design

We designed a fast event-related fMRI paradigm during which participants listened to pairs of color and action words. In each trial the two words were played one after the other with a stimulus onset asynchrony (SOA) of 2000 ms.

The inter-trial interval ranged between 4000 and 16000 ms. Participants were asked to judge the similarity of the two colors or the two actions from 1 to 5 (1: very different, 5: very similar). Responses were collected via an ergonomic hand-shaped response box with five keys (Resonance Technology Inc.). All participants used their right hand to provide responses (thumb = very different, pinky = very similar). Participants were told that they had about 4 seconds to provide a response after the onset of the second word of the pair and they were encouraged to use all the scale (1 to 5). Furthermore, the instruction was to judge the similarity of colors and actions based on their perceptual properties (avoiding reference to emotion, valence, or other non perceptual characteristics). Blind participants were told to judge color pairs on the basis of their knowledge about the perceptual similarity between colors.

Color and action words were presented in all possible within-category combinations (15 color pairs, 15 action pairs). Each pair was presented twice in each run, in the two possible orders (e.g., red-yellow, yellow-red). Thus, there were 60 trials in each run and the experiment consisted in 5 runs of 7 minutes. Stimuli were pseudorandomized using optseq2 to optimize the sequence of presentation of the different conditions. Three different optimized lists of trials were used across runs. List order was counterbalanced across subjects.

One early blind was excluded from the analyses because the subject answered to less than 70% of the trials throughout the experiment due to sleepiness. One run of one sighted subject was excluded from the analysis because of a technical error during the acquisition, and two other runs (one in a sighted subject, one in a blind subject) were excluded since the subject answered to less than 70% of the trials in that specific run.

### Conceptual similarity ratings

In order to perform the adaptation analysis, we divided the trials in similar pairs (e.g. red - orange) and different pairs (e.g. red - blue). We did so based on the participants’ subjective ratings. For each participant we took the average rating for each of the 15 word-pairs in the action and color categories. Then we automatically divided the 15 pairs in 5 intervals (4 quantiles) of nearly equal size. This subdivision was performed using the function *quantile*, in R (R Core Team, 2013), that divides a probability distribution into contiguous intervals of equal probabilities (i.e., 20%). The pairs in the first two intervals were the different pairs (low ratings of similarity), the pairs in the 3^rd^ interval were the medium pairs, and the pairs in the 4^th^ and 5^th^ intervals were the similar pairs (See fig 2B). However, in some cases, ratings distributions were slightly unbalanced, due to the tendency of some subjects to find more “very different” pairs than “very similar” pairs. In these cases (8 subjects for action ratings [3 EB]; 4 subjects for Color Ratings [1 EB]), the automatic split in 5 equal intervals was not possible. Thus, we set the boundary between the 2^nd^ and 3^rd^ interval at the ratings average (for that given subject), and set to the minimum (1 or 2, depending on the cases) the number of items in the 3^rd^ interval (not analyzed), in order to balance as much as possible the number of pairs in the Different and Similar groups. This procedure made so that in these special cases (as well as in all the others), the rating values of different pairs were always below the mean, and the values of similar pairs was always above the mean. Fig. S4, S5, S6, S7, in the supplementary information, show subject-specific rating distributions.

### MRI data acquisition

Brain images were acquired at the Neurological Institute Carlo Besta in Milano on a 3-Tesla scanner with a 32-channel head coil (Achieva TX; Philips Healthcare, Best, the Netherlands) and gradient echo planar imaging (EPI) sequences.

In the event-related experiment, we acquired 35 slices (voxel size 3 X 3 X 3.5) with no gap. The data in-plane matrix size were 64 X 64, field of view (FOV) 220mm X 220mm, time to repetition (TR)= 2 s, flip angle 90 degrees and time to echo (TE)= 30 ms. In all, 1210 whole-brain images were collected during the experimental sequence. The first 4 images of each run were excluded from the analysis for steady-state magnetization. Each participant performed 5 runs, with 242 volumes per run.

Anatomical data was acquired using a T1-weighted 3D-TFE sequence with the following parameters: 1 X 1 X 1 mm voxel size, 240 X 256 matrix size, 2.300 ms TR, 2.91 ms ET, 900 ms TI, 256 FoV, 160 slices.

### MRI data analysis

We analyzed the fMRI data using SPM12 (www.fil.ion.ucl.ac.uk/spm/software/spm12/) and Matlab R2014b (The MathWorks, Inc.).

#### Preprocessing

Preprocessing included slice timing correction of the functional time series (Sladky et al., 2011), realignment of functional time series, coregistration of functional and anatomical data, spatial normalization to an echoplanar imaging template conforming to the Montreal Neurological Institute (MNI) space, and spatial smoothing [Gaussian kernel, 6 mm full-width at half-maximum (FWHM)]. Serial autocorrelation, assuming a first-order autoregressive model, was estimated using the pooled active voxels with a restricted maximum likelihood procedure, and the estimates were used to whiten the data and design matrices.

#### Data analysis

Following preprocessing steps, the analysis of fMRI data, based on a mixed-effects model, was conducted in two serial steps accounting, respectively, for fixed and random effects. In all the analysis the regressors for the conditions of interest consisted of an event-related boxcar function convolved with the canonical hemodynamic response function according to a variable epoch model (Grinband, Wager, Lindquist, Ferrera, & Hirsch, 2008). Movement parameters derived from realignment of the functional volumes (translations in x, y, and z directions and rotations around x, y, and z axes), and a constant vector, were also included as covariates of no interest. We used a high-pass filter with a discrete cosine basis function and a cutoff period of 128 s to remove artifactual low-frequency trends.

#### Adaptation analysis

For each subject, the general linear model included 6 regressors corresponding to the 3 levels of similarity (different, medium, similar) in each condition (color, action). Color and Action pairs in the medium condition were modeled as regressors of no interest.

At the first level of analysis, linear contrasts tested for Repetition Suppression [Different > Similar] collapsing across categories (Action, Color). The same contrasts were then repeated within each category [Color Different > Color Similar; Action Different > Action Similar]. Finally, we tested for the Similarity by Category interactions, testing whether the adaptation was stronger in one category compared to the other (e.g., [Color Different > Color Similar] > [Action Different > Action Similar]).

These linear contrasts generated statistical parametric maps [SPM(T)]. The resulting contrast images were then further spatially smoothed (Gaussian kernel 5mm FWHM) and entered in a second-level analysis (RFX), corresponding to a random-effects model, accounting for inter-subject variance. One-sample t-tests were run on each group separately. Two-sample t-tests were then performed to compare these effects between groups (Blind vs Sighted).

#### Univariate analysis

For each subject, changes in regional brain responses were estimated through a general linear model including 2 regressors corresponding to the two categories Action and Color. The onset of each event was set at the beginning of the first word of the pair, the offset was determined by the subject response, thus included reaction time (Grinband et al., 2008). Linear contrasts tested for action-specific [Action > Color] and color-specific [Color > Action] BOLD activity.

These linear contrasts generated statistical parametric maps [SPM(T)]. The resulting contrast images were then further spatially smoothed (Gaussian kernel 5mm FWHM) and entered in a second-level analysis, corresponding to a random-effects model, accounting for inter-subject variance. One-sample t-tests were run on each group separately. Two-sample t-tests were then performed to compare these effects between groups (Blind vs Sighted) and to perform conjunction analyses to observe if the two groups presented similar activated networks for the two contrasts of interests.

#### Connectivity analysis

Psychophysiological interaction (PPI) analyses were computed to identify brain regions showing a significant change in the functional connectivity with seed regions (the right precuneus, the left pMTG and the rIPS) that showed a significant activation (p<.001, uncorrected) in the [(EB Conj. SC) X (Color > Action)] contrast, the [(EB Conj. SC) X (Action > Color)] contrast, and the [(SC > EB) X (Action > Color)] contrast respectively. In each individual, time series of activity (principal eigenvariate) were extracted from a 8 mm sphere centered on the nearest local maxima to the identified peaks in the second-level analysis (Note that centering the sphere on the peak itself does not change the ROI Analysis results, see Supplementary Information). New linear models were generated at the individual level, using three regressors. One regressor represented the psychological condition of interest (action or color trial). The second regressor was the physiological activity extracted in the reference area. The third regressor represented the interaction of interest between the first (psychological) and the second (physiological) regressor. The design matrix also included movement parameters and a constant vector as regressors of no interest. A significant PPI indicated a change in the regression coefficients between any reported brain area and the seed region, related to the experimental conditions (Color>Action or Action>Color). Next, the individual summary statistic images obtained at the first-level (fixed-effects) analysis were spatially smoothed (5 mm FWHM Gaussian kernel) and entered in a second-level (random-effects) analysis using a two-sample t test contrasting the two groups.

#### ROI definition

Occipital ROI for the PPI analyses were defined as following. Two peak-coordinates were taken from previous studies (Bedny et al. 2011; Kanjlia et al. 2016) showing the involvement of EB occipital areas in high-level functions such as language (left MOG [-36, −90, −1]) and mathematics (right MOG [33, −82, 9]). These areas also showed increased long range connectivity (in early blind) with frontal and parietal areas during rest (Bedny et al. 2011; Kanjlia et al. 2016; Liu et al. 2007; Collignon et al. 2013).

The V4 and V5 ROI were drawn from the literature, considering both perceptual localizers, as well as evidence from semantic/conceptual task. We selected 3 peak coordinates for area V5. The first [-47, −78, −2] from a highly-cited study contrasting the perception of visual motion vs static images (Dumoulin et al., 2000). The second [-44, −74, 2] from a study (Saygin, McCullough, Alac, & Emmorey, 2010) showing V5 sensitivity to motion sentences (e.g., “The wild horse crossed the barren field”). The third from a research on the on-line meta-analysis tool Neurosynth (http://neurosynth.org/) for the topic “action”. In Neurosynth, the area in the occipital cortex with the highest action-related activation was indeed V5 (peak coordinates: −50, −72, 2). To avoid ROI proliferation, we averaged these 3 peak-coordinates in order to obtain a single peak (average peak: −47, −75, 1).

As for V4, we selected the color-sensitive occipital ROI considering perceptual localizers, as well as evidence of color-specific activity from semantic/conceptual task. Fernandino et al. (Fernandino et al., 2015) reported a color sensitive area in the left posterior collateral sulcus (ColS; at the intersection between the Lingual and the Fusyform gyrus; MNI peak coordinates: −16, −71, −12) associated with color-related words. This peak is close to the posterior-V4 localization done by Beauchamps and colleagues (peak coordinates: −22, −82, −16) in a MRI version of the Farnsworth– Munsell 100-Hue Test (Beauchamp, Haxby, Jennings, & DeYoe, 1999). A search in neurosynth with the keyword “color” also highlighted a left posterior color-sensitive region along the ColS with peak coordinates [-24, −90, −10]. We averaged these 3 peaks to find the center of our region of interest (average peak: −21, −81, −13).

The posterior lateral-temporal cortex ROI (PLTC) was taken from 3 studies showing semantic repetition suppression in that area. Bedny and colleagues (Bedny, McGill, and Thompson-Schill 2008) observed increased neural adaptation in PLTC (peak coordinates: 57, −36, 21) for repeated words (fan - fan), when the words were presented in a similar context (summer – fan; ceiling - fan), compared to when different context triggered different meanings (e.g., admirer – fan; ceiling – fan). This result conceptually replicated previous studies (Kotz, Cappa, Cramon, & Friederici, 2002; Wible et al., 2006) showing semantic adaptation in the bilateral PLTC for related (e.g., dog - cat) vs unrelated (e.g., dog - apple) word pairs (peak coordinates: −42, −27, 9 and −51, −22, 8). These 3 peaks were averaged to find the center of our region of interest in both hemispheres (average peak: ±50, −28, 13).

## Statistical analysis

At the whole brain level, statistical inference was made at a corrected cluster level of P < 0.05 FWE (with a standard voxel-level threshold of P < 0.001 uncorrected) and a minimum cluster-size of 50 voxels. ROI analysis based on Small Volume Correction were thresholded at p<0.05 FEW at the voxel level.

All ROI analyses were performed using Small Volume Correction using spheres with a 10mm radius centered around the ROI peak coordinates (see previous session). Within the ROI, results were considered significant at a threshold of p<0.05, FEW-corrected. Here, and throughout the paper, brain coordinates are reported in MNI space.

Behavioral data, analysis code and t-maps from the main contrasts will be made available on-line (www.biorxiv.org/content/early/2018/08/23/384552). Row fMRI images will be made available upon request, following agreements with our ethical board committee.

## Results

### Within-category similarity is encoded in occipital areas in the sighted but not in the blind

The rationale behind adaptation analyses was that the direct contrast between pairs with high versus low perceptual differences will display neural adaptation (Barron et al., 2016; Grill-Spector, Henson, & Martin, 2006; Wheatley, Weisberg, Beauchamp, & Martin, 2005) therefore probing regions that are specifically sensitive to the *perceptual distance* between concepts.

*Behavioral analysis*: Similarity ratings were highly correlated between sighted and blind, both for action (r= .99) and color concepts (r= .93; Fig. 1A). In order to perform the adaptation analysis we divided the trials in similar pairs (e.g. red - orange) and different pairs (e.g. red - blue), based on each participant’ subjective ratings. Rating distributions for each subject and category (color, action) were divided in 5 intervals with a similar number of items (see Method session for details). Stimulus-pairs in the first two intervals were labeled as different (low similarity ratings), the 3^rd^ interval contained medium pairs, and the 4^th^-5^th^ intervals similar pairs (high similarity ratings; Fig 1B). Overall, the average number of “different” trials was slightly larger than the “similar” ones (126 vs 115; F(1,33)=8.41, p=0.007, *η*^2^=0.20; Fig. 1C). However, there was no similarity by group interaction (F(1,33)=0.18, p=0.67, *η*^2^=0.004), indicating that this unbalance (that reflected personal judgments of similarity) was the same across SC and EB (fig. 1C). An analysis of reaction times showed that Medium pairs (not analyzed in fMRI) had on average longer latencies than Similar and Different ones (Main Effect of Similarity: F(2,66)=21.07, p<0.001, *η*^2^=0.38). This was expected since pairs that are neither similar nor different would require longer and more difficult judgments. Crucially, there was no difference in reaction times between different (Mean=1.80 sec, SD=0.39) and similar pairs (Mean=1.79 sec, SD=0.37; F(1,33)=0.09, p=0.76, *η*^2^=0.003), and no interaction between Similarity and Group (F(1,33)=0.04, p=0.84, *η*^2^=0.001; Fig 1D).

**Figure 1.**
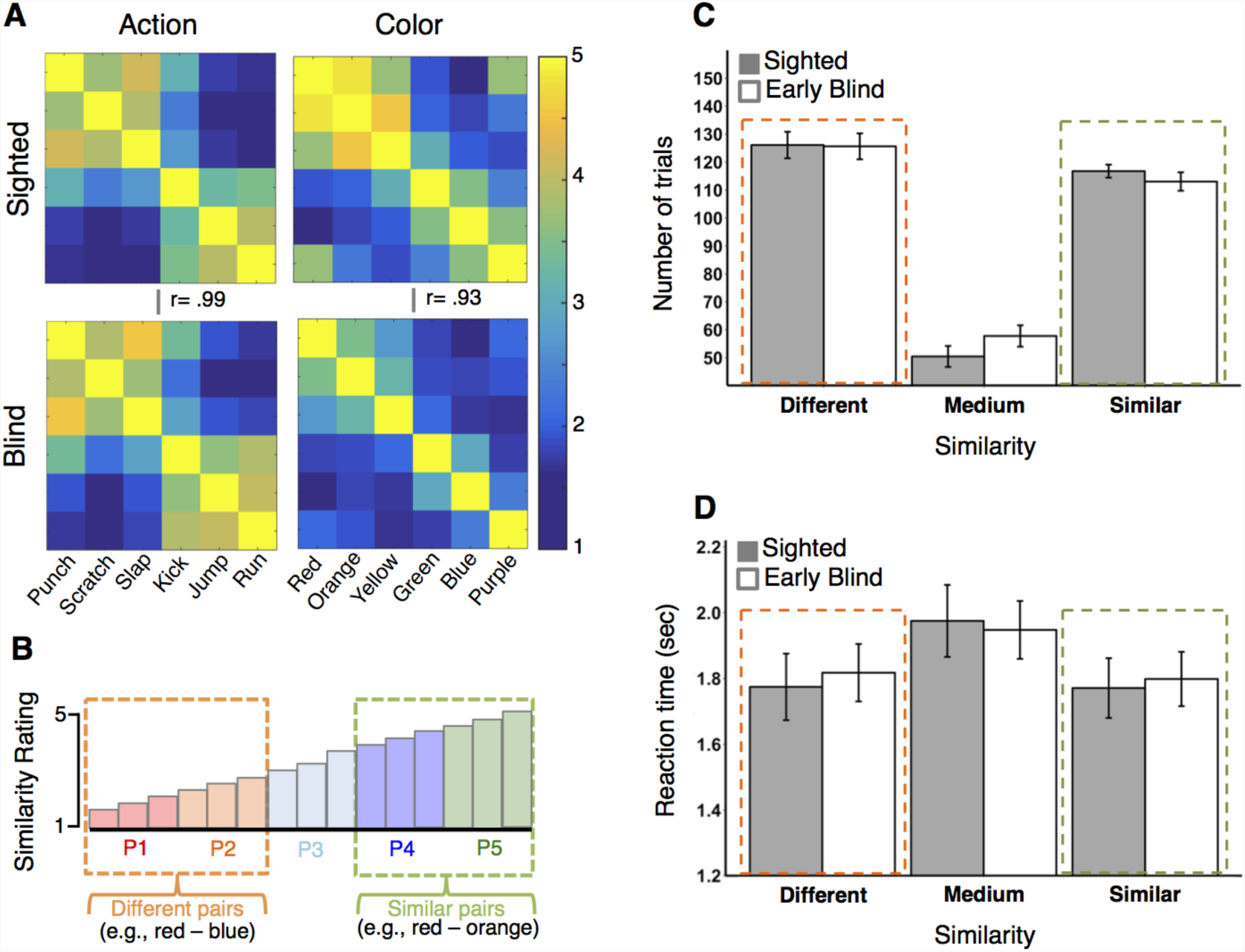
Adaptation, behavioral analysis. (A) Similarity judgments were highly correlated across groups both for actions and color; (B) Conceptual schema of the division of word pairs in “different” and “similar” based on subjective similarity ratings; (C) Barplot depicting the average number of items in the “different”, “medium” and “similar” categories. The number of items in the “different” and “similar” categories is very similar across groups (number of trials ± SEM); (D) Barplot depicting the average reaction time in the “different”, “medium”, and “similar” categories. The average RTs of the “different” and “similar” categories is very similar across groups (seconds ± SEM).

*fMRI analysis*: to find brain areas that showed adaptation based on conceptual similarity, we looked at the contrast Different Pairs > Similar Pairs, with Medium pairs as a regressor of no interest. Action and color pairs were considered together since, at the whole brain level, we did not find a significant higher-order interaction between Similarity (different, similar) and Category (see the method section for analysis details). In the sighted, similar concepts led to repetition suppression in several occipital areas (See Fig. 2A), with a significant cluster in and around the left lingual gyrus (peak coordinates: −24, −70, −7). In the blind, instead, adaptation emerged in language-related areas with significant clusters along the middle and superior temporal gyrus, bilaterally (Peak coordinates RH: 57, −28, 8; LH: −60, −10, −7) and in the right precentral gyrus (Peak coordinates: 27, −25, 56; Fig. 2B). Importantly, no adaptation in posterior occipital areas was observed in the blind.

**Figure 2.**
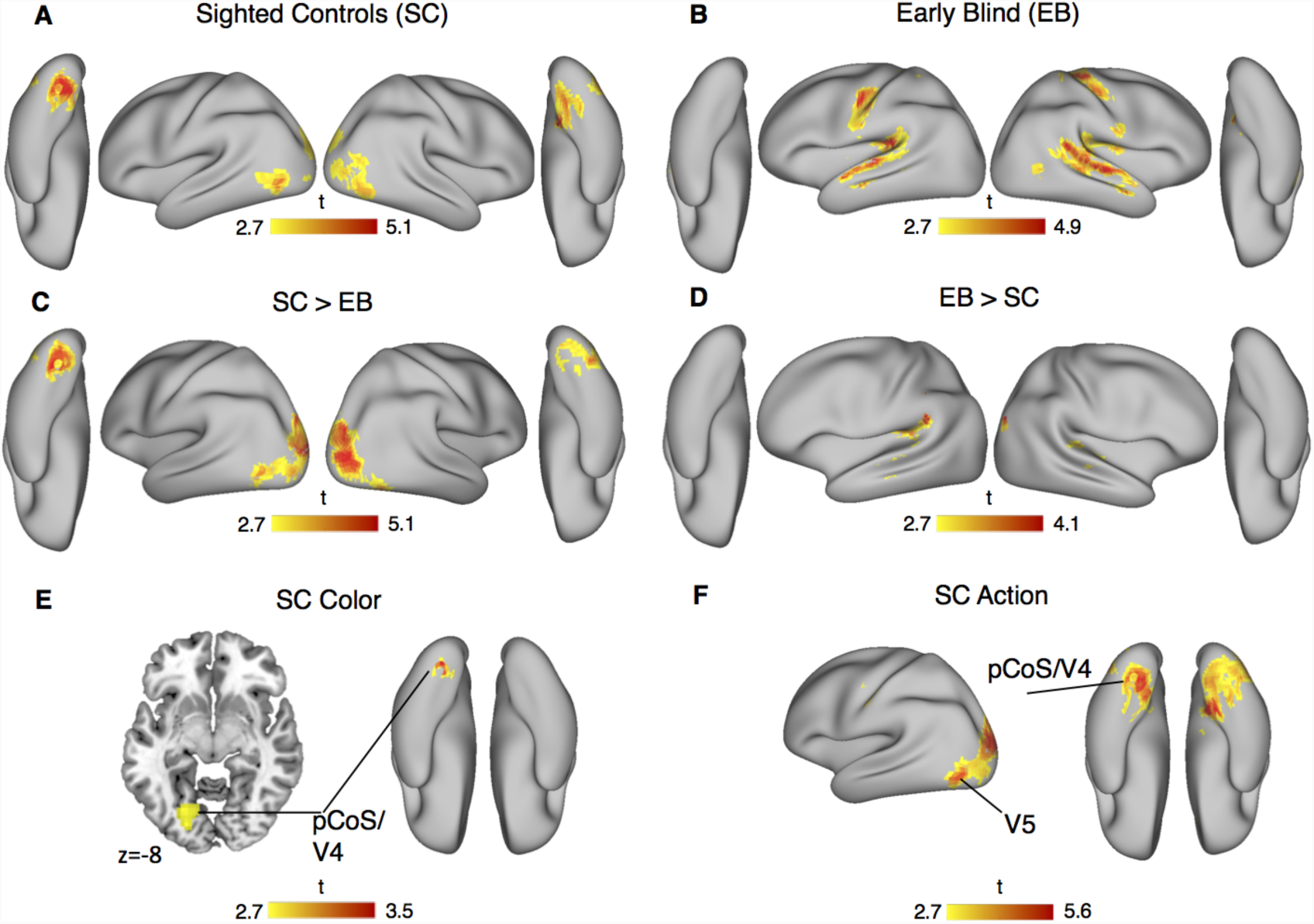
Adaptation, fMRI results. Regional BOLD responses are rendered over Conte-69 average midthickness surfaces. (A) Suprathreshold clusters showing neural adaptation for similar word pairs in the occipital cortices of sighted participants; (B) Suprathreshold clusters showing neural adaptation for similar word pairs in the temporal and somatosensory-motor cortices of early blind participants; (C) Suprathreshold clusters showing neural adaptation for similar word pairs in sighted compared to blind, and (D) blind compared to sighted; (E) Suprathreshold cluster showing neural adaptation for similar color-word pairs in the sighted, with regional activity precisely localized in the left PCoS/V4 (see Fernandino et al. 2015 for a very similar result with a multivariate design); (F) Suprathreshold clusters showing neural adaptation for similar action-word pairs in the sighted, with regional activity spread in different occipital areas including the left PCoS/V4 and the left V5. Cluster threshold at P<0.005 uncorrected, for illustration only.

A comparison between groups showed greater adaptation in occipital cortices for sighted compared to blind (Fig. 2C), with peaks in the left superior occipital gyrus (−24, −91, 26), the left lingual gyrus (−24, −70, −7) and the right middle occipital gyrus (27, −85, 11). The contrast Blind > Sighted showed increased adaptation in posterior lateral temporal cortices (PLTC) bilaterally (Fig. 2D). Planned ROI analysis in PLTC, a region that consistently show repetition suppression for semantic similarity (Bedny, McGill, and Thompson-Schill 2008; Wible et al. 2006; Kotz et al. 2002), revealed a significantly greater adaptation for similar concepts in blind more than sighted (Conceptual Similarity by Group interaction; lPLTC= −45 −31 20, t(33)=3.23, P*=* 0.035; rPLTC= 45 −28 11, t(33)=3.41, P*=* 0.024).

Finally, we performed planned ROI analysis in the color-sensitive region at the posterior banks of the collateral sulcus (PCoS) corresponding to the V4-complex (Beauchamp et al., 1999; Fernandino et al., 2015) and the motion sensitive region V5 (Dumoulin et al., 2000). In area PCoS-V4 we found greater adaptation both for color and action in sighted compared to blind (Conceptual Similarity by Group interaction; peak= −24 −73 −10, t(33)=4.26, P*=* 0.004; Fig 2 E-F). In contrast, the analysis in V5 showed that repetition suppression was specific for action concepts in the sighted and no adaptation was observed in the blind (Conceptual Similarity by Group by Category interaction; Peak: −51 −76 8, t(33)=3.29, P= 0.037; Fig 2F).

### Brain regions active in sighted and blind when contrasting action and color concepts

Subsequently we ran classic univariate analysis, comparing items across categories independently of their similarity, to find category-specific activations across sighted and blind. In these analyses, the two words in each pair were considered as a single trial.

*Behavioral analysis*: Reaction times analysis using a mixed ANOVA, with Category (action, color) as within-subject factor and Group (sighted, blind) as between-subjects factor, showed no difference between categories (F(1,33)=2.37, p>0.05, *η*^2^= 0.07), between groups (F(1,33)=0.074, p>0.05, *η* ^2^=0.002) and no Category by Group interaction (F(1,33)=0.69, p>0.05, *η*^2^=0.02).*fMRI analysis*: The contrast Action > Color did not reveal any significant difference between groups, suggesting a comparable categorical representation of action concepts, across sighted and blind (see fig. S2 for details). Indeed, a conjunction analysis between groups showed a common significant activation in the left posterior middle temporal gyrus (lpMTG; Peak= −54, −61, 5; Fig 3A).

**Figure 3.**
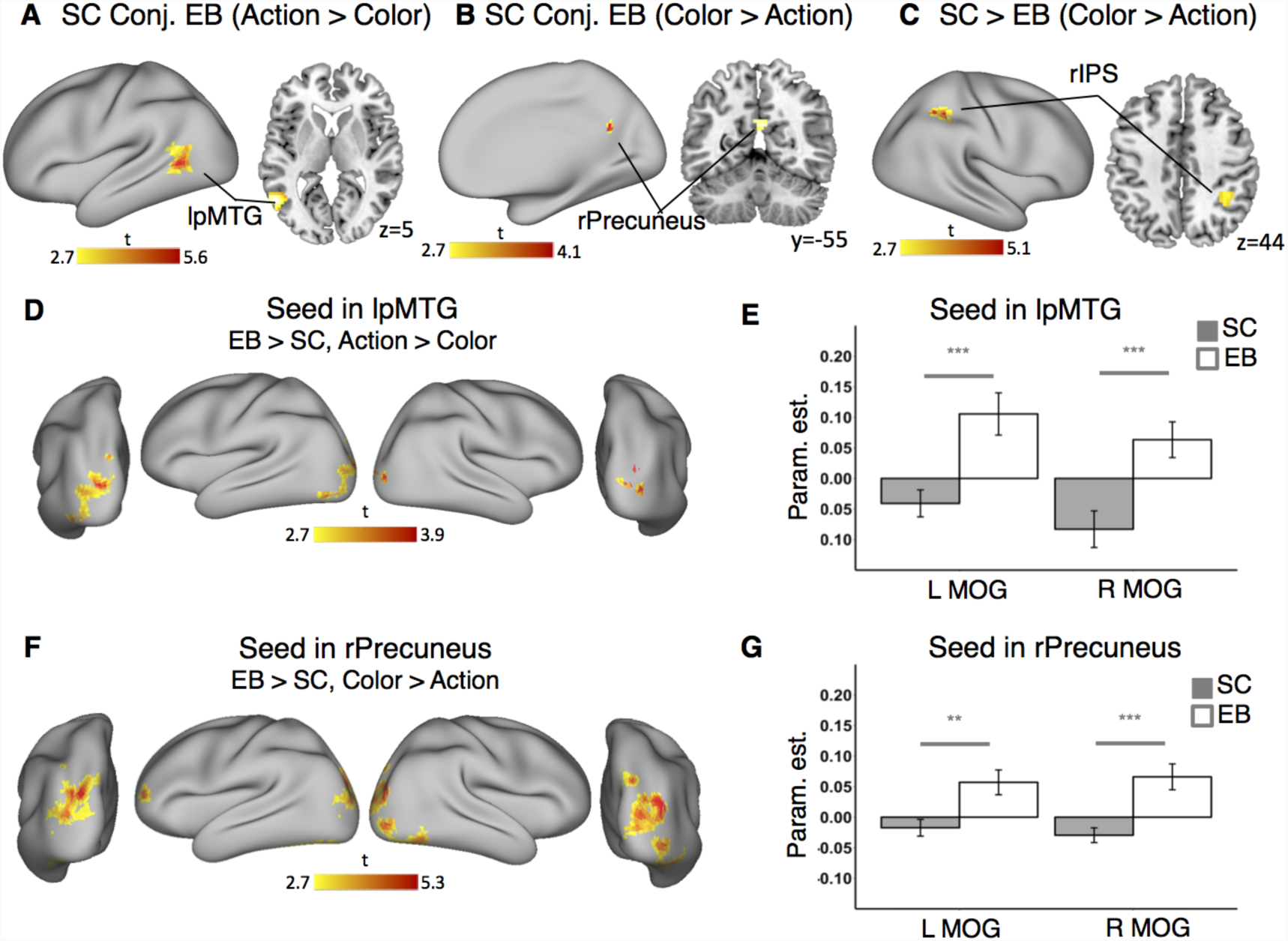
Contrasts between categories and psychophysiological interactions. Regional BOLD responses are rendered over Conte-69 average midthickness surfaces. (A) Suprathreshold cluster (P<0.05 FEW corrected) showing common activity in the lpMTG for the contrast Action > Color in both sighted and early blind (conj., conjunction analysis); (B) Suprathreshold cluster (P<0.001 uncorrected; for illustrative purposes only) showing common activity in the precuneus for the contrast Color > Action in both sighted and early blind (conj., conjunction analysis); (C) Suprathreshold cluster (P<0.05 FEW corrected) showing greater activity in the rIPS, in sighted compared to early blind, for the contrast Color > Action; (D) Suprathreshold clusters (P<0.005 uncorrected; for illustrative purposes only) showing greater connectivity in the occipital areas of early blind people with the lpMTG (PPI for the contrast Action > Color); (E) Barplot (for illustrative purposes only) showing beta weights derived from PPI analysis, with seed in lpMTG, in sighted (gray) and blind (white), in the right and left middle occipital gyrus (MOG) for the contrast Action > Color (arbitrary unit ± SEM); (F) Suprathreshold clusters (P<0.005 uncorrected; for illustrative purposes only) showing greater connectivity in the occipital areas of early blind people with the precuneus (PPI for the contrast Color > Action); (G) Barplot (for illustrative purposes only) showing beta weights derived from PPI analysis, with seed in precuneus, in sighted (gray) and blind (white), in the right and left middle occipital gyrus (MOG) for the contrast Color > Action (arbitrary unit ± SEM).

On the other hand, a conjunction analysis between groups for Color > Action did not reveal any common activation between sighted and blind after correction for multiple comparisons at the whole brain level. However, displaying the conjunction results at a more lenient threshold (p<.001 uncorrected; Fig 3C), we could notice a unique common activity for color concepts in the right precuneus (peak= 6, −55, 26). Accordingly, analysis within groups showed a significant precuneus activity in the blind (peak= 6, −52, 20, p= .04) and a marginally significant activity in the sighted (peak= 0, −61, 29, p= .06), with no significant difference between groups (Table S1; Fig. S1).

Further analysis for the contrast Color > Action revealed a cluster in the right parietal cortex, in and around the right intraparietal sulcus (rIPS), showing higher activity for color concepts in sighted compared to blind (peak= 33, −43, 35; Fig. 3B).

Altogether these results show similar patterns of activity during conceptual processing in sighted and blind, when categorical preferences are investigated (independently of perceptual similarity). As in previous results (Bedny et al., 2012; Uta Noppeney et al., 2003; van den Hurk et al., 2017) such common activities are found outside the posterior occipital cortex, in areas that are considered to be highly polymodal such as the precuneus and the lpMTG (Binder, Desai, Graves, & Conant, 2009). Interestingly, even at this level of comparison we could find an effect of visual deprivation, concerning color knowledge, that seem to involve the right IPS more in sighted than blind.

### Psychophysiological interactions: The precuneus and lpMTG display similar category selectivity across sighted and blind but show different connectivity profiles

Finally, we relied on Psychophysiological Interaction (PPI) analysis (Friston et al., 1997; O’Reilly, Woolrich, Behrens, Smith, & Johansen-berg, 2012) to test whether regions that showed similar categorical preference across groups (lpMTG for action > color, and the precuneus for color > action) also maintain a similar connectivity profile in both groups. In particular, we tested the hypothesis that posterior occipital areas in the blind could be recruited during conceptual retrieval, and connect with conceptual hubs (Binder et al., 2009) such as the lpMTG and the Precuneus, as a consequence of neural reorganization (Amedi, Raz, Pianka, Malach, & Zohary, 2003; Burton, 2003). With this aim, we selected two ROIs in the left and right middle occipital gyrus (MOG) that are recruited in early blind during high-level conceptual tasks such as language processing and math (Kanjlia et al. 2016; Bedny et al. 2011); and show increased long range connectivity, in early blind, with extra-occipital regions (e.g., frontal, parietal and ventral temporal cortices) during resting state (Bedny et al. 2011; Kanjlia et al. 2016; Liu et al. 2007; Collignon et al. 2013) and task-based (PPI) analysis (Noppeney, Friston, and Price 2003).

PPI with seed in the lpMTG revealed an increase of action-selective functional connectivity in both occipital ROIs of blind people compared to their sighted counterpart (lMOG: t(33)= 3.59, p= 0.02; rMOG: t(33)=3.46, p=0.026; Fig 1D & 1E). Similarly, PPI with seed in the precuneus revealed an increase of color-selective functional connectivity in the occipital cortex of blind compared to sighted participants (lMOG: t(33)= 3.12, p= 0.054; rMOG: t(33)=4.09, p=0.007; Fig 1F and 1G). Albeit showing a similar activity profile in sighted and blind during conceptual processing, the lpMTG and the precuneus showed a different connectivity profile as a function of early visual deprivation. Such increase in task-based connectivity suggests that occipital areas in early blind, albeit not coding for perceptual similarity as in the sighted, are however active during conceptual retrieval and can be flexibly recruited in interaction with conceptual hubs such as the precuneus and the pMTG.

## Discussion

Embodied approaches to conceptual knowledge suggest that concepts are grounded in our sensory and motor experience of the world (Barsalou, 1999; Binder & Desai, 2011). A straightforward hypothesis emerging from these theories is that people that perceive the world in a different way should also have different conceptual representations (Casasanto, 2011).

In our study we tested this hypothesis by characterizing the brain activity of sighted and early blind individuals while they rated the perceptual similarity of action and color concepts in fMRI. In particular, we investigated which brain regions encode the perceptual similarity of retrieved concepts using an adaptation paradigm. Results in the sighted group showed that word-pairs referring to similar colors or actions induced repetition suppression in several posterior occipital regions, including areas V3, V4 and V5. In striking contrast, early blind participants did not show repetition suppression in posterior occipital areas but instead showed a greater adaptation for concept similarity in language-related regions, disclosing a different neurobiology of concepts as a function of visual deprivation.

Posterior occipital regions are known to encode visual features in the sighted and to be sensitive to visual similarity independently of categorical membership (Bracci and Op de Beeck 2016; Connolly et al. 2012; Kriegeskorte et al. 2008; Naselaris et al. 2009). Our data corroborate the hypothesis that these regions are also involved in conceptual retrieval (Borghesani et al., 2016; Fernandino et al., 2015) supporting information related to the visual appearance of objects and events, that is not available to blind people. This result is crucial to support the hypothesis that conceptual retrieval consists, in part, on replay of our perceptual experience (Barsalou, 1999; Harnad, 1990).

Since posterior occipital regions represent visual features also during conceptual retrieval (Borghesani et al., 2016; Fernandino et al., 2015; Mitchell et al., 2008), it is therefore expected, as observed in our study, that sighted and blind should show different activity in these regions due to the lack of visual qualia in the blind population. Our results are in line with a previous study showing that relatively anterior regions such as the pMTG and part of the VOTC show a similar functional and connectivity fingerprint in sighted and blind (Wang et al., 2015), whereas this similarity decreases strikingly in more posterior occipital regions (i.e., approximately behind the conventional line that separate the temporal from the occipital lobe; Wang et al. 2015). Actually, the posterior occipital regions showing higher conceptual adaptation in sighted compared to blind in our study tightly overlap with regions showing the lowest functional and connectivity similarity between sighted and blind in Wang et al. study (Wang et al. 2015; Fig. S2).

Reduced adaptation in the occipital cortex in our blind participant (compared to sighted) coincides with stronger adaptation in the posterior lateral-temporal cortices (PLTC). Several studies have found that conceptually similar (e.g., dog - wolf) or semantically associated (e.g., dog - leash) words can lead to repetition suppression in PLTC (Wible et al. 2006; Kotz et al. 2002; Bedny, McGill, and Thompson-Schill 2008). Although it is still unclear what level of conceptual knowledge is represented in that region (Bedny, McGill, and Thompson-Schill 2008), there is some agreement that the PLTC stores auditory representations of words, that are connected to distributed semantic representations in the brain (Hickok and Poeppel 2007; Gow 2012; Bedny, McGill, and Thompson-Schill 2008; Mirman et al. 2015). In this framework, the PLTC may work at the interface between wordforms and semantic knowledge (Gow, 2012; Hickok & Poeppel, 2007), and a greater activity in the blind can index a larger use of verbal knowledge in this population (Cattaneo et al., 2008; Crollen et al., 2014).

Directly contrasting different categories (Action Vs Color), independently of within-category similarity, showed instead some commonalities in brain activity across the two groups (Bedny et al. 2012; Noppeney, Friston, and Price 2003; Peelen et al. 2014). Both sighted and blind engaged the lpMTG during action processing and the Precuneus during color processing. These regions are located outside of the visual cortex, in polymodal areas, that typically display category-specificity in a format that is at least partially independent from perceptual appearance (Bracci & Op de Beeck, 2016; Peelen & Downing, 2017) and largely resilient to visual deprivation (Bedny et al., 2012; Uta Noppeney et al., 2003; Wang et al., 2015).

Interestingly, though, psychophysiological interactions (PPI) showed that early visual deprivation changes the connectivity profile of these regions, increasing their functional coupling with occipital regions in the blind. These results suggest that the occipital cortex in early blind is re-organized to extend its integration into conceptual selective networks, highlighting further how visual deprivation impact on the neurobiology of conceptual knowledge. Previous studies have shown that occipital areas in the early blind are recruited for high-level conceptual tasks such as language processing, semantic retrieval and math (Bedny et al. 2011; Kanjlia et al. 2016; Burton, Diamond, and McDermott 2003; Van Ackeren et al. 2018; Crollen et al. 2018; See also Fig. S3 showing general higher activity in the blind occipital cortex during the auditory presentation of words), and that they increase long-range connectivity with frontal and parietal cortices during rest (Bedny et al. 2011; Kanjlia et al. 2016; Liu et al. 2007; Collignon et al. 2013) and inferior temporal cortices during semantic judgments (Noppeney, Friston, and Price 2003). Notably, graph-theoretic metrics of regional cortical thickness covariance found that language and visual regions showed a pattern of merging into shared modules in the blind but not in sighted (Hasson, Andric, Atilgan, & Collignon, 2016). Extending those previous studies, we show here that early blinds seem to rely on enhanced connectivity between occipital cortices and temporo-parietal conceptual hubs during conceptual processing. Albeit our data remain correlational, they suggest that EBs’ occipital regions are not activated independently of other “classic” regions involved in conceptual retrieval, but they instead work in concert.

These results suggest that occipital cortices are involved in conceptual retrieval in both sighted and blind, but with different functions and probably at different levels of representation. Occipital areas may support sensorimotor simulations of visual features during conceptual retrieval in the sighted, showing adaptation for concepts that refer to visually similar objects or events; on the other hand, in the blind, occipital cortices do not encode perceptual similarity, but may be re-organized to engage in more general processes related to conceptual retrieval (albeit these processes need to be better specified; see for instance, Bedny 2017; Van Ackeren et al. 2018).

Outside the posterior occipital cortex, we found that the posterior portion of the right IPS showed a stronger preference for color trials in the sighted compared to the blind. The IPS is known to be involved in the perception of color (Beauchamp et al., 1999; Cheadle & Zeki, 2014; Zeki & Stutters, 2013) as well as other visual features (Grill-Spector, 2003; Swisher, Halko, Merabet, McMains, & Somers, 2007; Xu, 2007), and its anatomical position make it a good candidate to work at the interface between perceptual and conceptual representations (Cheadle & Zeki, 2014). In particular, it seems to be crucial for retrieving perceptually-based categorical knowledge (Cheadle & Zeki, 2014; Zeki & Stutters, 2013). Indeed, the peak of color-specific activity that we found (peak coordinates: 33, −43, 35) is very close to the color-specific rIPS area found by Cheadle & Zeki (2014; peak coordinates: 30, −39, 45). The lack of visual input in blind people prevents the formation of perceptually-driven color representations, which may limit the contribution of the IPS during the retrieval of color knowledge. This is not the case for action representation, for which perceptual knowledge can be more easily compensated by other senses (e.g., touch, audition) in the blind. The distinct impact of blindness on the broader categorical responses to action and colors again highlight how visual experience may selectively shape the neurobiology of conceptual representations, even in putatively multimodal areas of the brain.

From a broader theoretical point of view, the results of this study are in line with a hierarchical model of conceptual representations based on progressive levels of abstraction (Barsalou, 2016; Binder, 2016; Binder et al., 2016; Fernandino et al., 2015; A. Martin, 2015). At the top of the hierarchy, multimodal representations may co-exist with purely symbolic ones organized in a linguistic/propositional code (Mahon & Caramazza, 2008). This level of representation can account for the obvious fact that congenitally blind people can think about colors and their perceptual properties (Barilari, de Heering, Crollen, Collignon, & Bottini, 2018; Marmor, 1978) although they cannot simulate vision. On the other hand, modality-specific simulation in sensory areas (e.g. visual, auditory, somatosensory), as the one highlighted by our adaptation analysis, may become central in deeper and more deliberate stages of conceptual processing, providing situated, specific and sometimes imaginistic representations, and eventually supporting the subjective experience of knowing providing the phenomenological qualia of conscious thinking (Binder et al., 2016; Koch, Massimini, Boly, & Tononi, 2016).

## Supporting information

## Acknowledgement

This work was supported by a European Research Council starting grant (MADVIS grant #337573) attributed to OC. OC is a research associate at the Fond National de Recherche Scientifique de Belgique (FRS-FNRS). We wish to extend our gratitude to the Michela Picchetti, Mattia Verri and Alberto Redolfi for the technical support during fMRI acquisition. We are also extremely thankful to our blind participants and the Unione Ciechi e Ipovedenti in Trento, Milano, Savona, Trieste, and the Blind Institute of Milano. We also thank Yanchao Bi and Xiaoying Wang for sharing brain maps from their previously published data.

