## Supplementary material for "Different neural networks for conceptual retrieval in sighted and blind"

### Supplementary Information

**Table S1. Demographic information of sighted and blind participants.**

| Sight Status | Age | Gender | Age of formal education | Sight Status | Age | Gender | Age of formal education |
|---|---|---|---|---|---|---|---|
| EB01 | 32 | M | 18 | SC01 | 37 | M | 18 |
| EB02 | 52 | F | 18 | SC02 | 46 | F | 18 |
| EB03 | 30 | M | 13 | SC03 | 34 | M | 18 |
| EB04 | 29 | F | 13 | SC04 | 26 | F | 16 |
| EB05 | 36 | F | 18 | SC05 | 33 | F | 18 |
| EB06 | 27 | F | 18 | SC06 | 27 | F | 18 |
| EB07 | 39 | M | 18 | SC07 | 40 | M | 18 |
| EB08 | 48 | M | 8 | SC08 | 51 | M | 8 |
| EB09 | 42 | M | 13 | SC09 | 41 | M | 18 |
| EB10 | 36 | M | 18 | SC10 | 40 | M | 18 |
| EB11 | 49 | M | 18 | SC11 | 43 | M | 18 |
| EB12 | 36 | F | 18 | SC12 | 39 | F | 18 |
| EB13 | 45 | M | 18 | SC13 | 42 | M | 18 |
| EB14 | 25 | F | 13 | SC14 | 25 | F | 16 |
| EB15 | 29 | F | 16 | SC15 | 30 | F | 18 |
| EB16 | 30 | M | 13 | SC16 | 26 | M | 13 |
| EB17 | 28 | F | 18 | SC17 | 31 | F | 18 |
| EB18 | 45 | M | 8 | SC18 | 39 | M | 13 |
| M | *36,56* | | *15,39* | | *36,11* | | *16,67* |
| SD | *8,48* | | *3,48* | | *7,52* | | *2,72* |

*Note.* M = male; F = female; EB=early blind; SC=sighted control; M=mean; SD=standard deviation.

**Table S2. Early blind participants, additional information**

| Subject | Handedness | Age onset blindness | Cause of blindness |
|---|---|---|---|
| EB01 | 32 | 0 | Optic nerve Hypoplasia |
| EB02 | 52 | 1 | Retinoblastoma |
| EB03 | 30 | 0 | Congenital retinal dystrophy |
| EB04 | 29 | 2 | Retinoblastoma |
| EB05 | 36 | 0 | Congenital Microphtalmia |
| EB06 | 27 | 0 | Congenital Microphtalmia |
| EB07 | 39 | 0 | Retrolental fibroplasia |
| EB08 | 48 | 0 | Optic nerve atrophy |
| EB09 | 42 | 0 | Congenital retinal dystrophy |
| EB10 | 36 | 0 | Retrolental fibroplasia |
| EB11 | 49 | 0 | Retrolental fibroplasia |
| EB12 | 36 | 0 | Retrolental fibroplasia |
| EB13 | 45 | 0 | Retrolental fibroplasia |
| EB14 | 25 | 0 | Leber's congenital amaurosis |
| EB15 | 29 | 3 | Retinitis pigmentosa |
| EB16 | 30 | 0 | Retrolental fibroplasia |
| EB17 | 28 | 0 | Agenesis of the optic nerve |
| EB18 | 45 | 0 | Glaucoma |

**Table S3. Regional responses for adaptation analysis (repetition suppression), across sighted and early blind.**

| Area | k | Xmm | Ymm | Zmm | *Z* | $P_{FWE}$ |
|---|---|---|---|---|---|---|
| Sighted, different > similar | | | | | | |
| L lingual gyrus | 180 | -24 | -70 | -7 | 4.33 | .013 |
| | S.C. | -15 | -70 | -10 | 4.32 | |
| Blind, different > similar | | | | | | |
| L middle temporal gyrus | 119 | -60 | -10 | -7 | 4.20 | .049 |
| | S.C. | -63 | -22 | 2 | 3.90 | |
| L superior temporal gyrus | S.C. | -51 | -13 | -1 | 3.39 | |
| R superior temporal gyrus | 323 | 57 | -28 | 8 | 4.23 | .001 |
| | S.C. | 48 | -37 | 20 | 3.92 | |
| | S.C. | 54 | -22 | 2 | 3.92 | |
| L Postcentral gyrus | 105 | -48 | -13 | 38 | 4.12 | .07‡ |
| R Precentral gyrus | 121 | 27 | -25 | 56 | 3.71 | .02 |
| | S.C. | 36 | -22 | 62 | 3.70 | |
| | S.C. | 36 | -16 | 44 | 3.45 | |
| Sighted > Blind, diff. > similar | | | | | | |
| L superior occipital gyrus | 141 | -24 | -91 | 26 | 4.35 | .03 |
| | S.C. | -9 | -97 | 20 | 3.77 | |
| L middle occipital gyrus | S.C. | -15 | -100 | 14 | 3.75 | |
| R middle occipital gyrus | 250 | 27 | -85 | 11 | 3.97 | .003 |
| | S.C. | 36 | -79 | 2 | 3.67 | |
| R superior occipital gyrus | S.C. | 27 | -79 | 23 | 3.79 | |
| L lingual gyrus | 165 | -24 | -70 | -7 | 3.91 | .017 |
| L middle occipital gyrus | S.C. | -39 | -73 | 2 | 3.48 | |
| L middle occipital gyrus | S.C. | -27 | -94 | 2 | 3.25 | |
| Blind > Sighted, diff. > similar | | | | | | |
| L superior temporal gyrus | | -45 | -31 | 20 | 2.99 | .035^ |
| R superior temporal gyrus | | 45 | -28 | 11 | 3.13 | .024^ |

Significance corrections are reported at the cluster level, unless otherwise specified; cluster size threshold = 50. FWE, family-wise error; L, left; R, right; S.C., same cluster. ^Brain activation significant after FWE voxel correction over a small spherical volume (10-mm radius) at peak coordinates for right/left PLTC. ‡ Indicates marginally significant clusters ($P_{fwe}<0.1$)

**Table S4. Regional responses for the comparison between action and color trials across sighted and early blind.**

| Area | k | Xmm | Ymm | Zmm | *Z* | $P_{FWE}$ |
|---|---|---|---|---|---|---|
| Sighted, action > color | | | | | | |
| L middle temporal gyrus | 1134 | -57 | -61 | 8 | 5.80 | <.001 |
| L inferior frontal gyrus | S.C. | -48 | 29 | -1 | 5.31 | |
| L middle temporal gyrus | S.C. | -60 | -43 | 29 | 5.15 | |
| R middle temporal gyrus | 474 | 60 | -10 | -7 | 5.14 | <.001 |
| | S.C. | 57 | 2 | -13 | 4.96 | |
| R superior temporal gyrus | S.C. | 51 | -28 | -1 | 4.74 | |
| Sighted, color > action | | | | | | |
| R orbital gyrus | 65 | 27 | 35 | -13 | 4.98 | .009* |
| L orbital gyrus | 71 | -30 | 35 | -13 | 4.90 | .012* |
| precuneus | 91 | 0 | -61 | 29 | 4.11 | .06 ‡ |
| Blind, action > color | | | | | | |
| L middle temporal gyrus | 315 | -54 | -61 | 5 | 4.69 | <.001 |
| | S.C. | -42 | -67 | 17 | 4.46 | |
| | S.C. | -45 | -52 | 14 | 3.49 | |
| R calcarine/post. cingulate | 325 | 9 | -67 | 8 | 4.63 | <.001 |
| | S.C. | 0 | -76 | 11 | 4.07 | |
| L calcarine | S.C. | -9 | -82 | 11 | 4.07 | |
| R middle temporal gyrus | 203 | 57 | -55 | 5 | 4.40 | .003 |
| | S.C. | 42 | -55 | 8 | 3.94 | |
| | S.C. | 63 | -43 | 8 | 3.63 | |
| R inferior frontal gyrus | 256 | 42 | 20 | 5 | 4.21 | <.001 |
| R superior temporal pole | S.C. | 48 | 11 | -19 | 3.97 | |
| R inferior frontal gyrus | S.C. | 51 | 17 | -1 | 3.80 | |
| Blind, color > action | | | | | | |
| R precuneus | 109 | 6 | -52 | 20 | 4.72 | .036 |
| L medial frontal gyrus | 174 | -9 | 59 | -7 | 4.20 | .008 |
| | S.C. | 6 | 53 | -10 | 3.85 | |
| Sighted ∩ Bind, action > color | | | | | | |
| L middle temporal gyrus | 155 | -54 | -61 | 5 | 4.69 | .01 |
| Sighted > Blind, color > action | | | | | | |
| R intraparietal sulcus | 146 | 33 | -43 | 35 | 4.33 | .013 |

Significance corrections are reported at the cluster level, unless otherwise specified; cluster size threshold = 50. FWE, family-wise error; L, left; R, right; S.C., same cluster.

*Brain activations significant after FWE correction at voxel-level over the whole brain. ‡ Indicates marginally significant clusters ($P_{fwe}$<0.1)

**Table S5. Regional responses for psychophysiological interactions with seeds in lpMTG and Precuneus, across sighted and early blind.**

| Area | k | Xmm | Ymm | Zmm | *Z* | $P_{FWE}$ |
|---|---|---|---|---|---|---|
| Seed in lpMTG | | | | | | |
| Sighted | | | | | | |
| L inferior parietal cortex | 174 | -45 | -40 | 41 | 4.12 | .006 |
| | S.C. | -36 | -52 | 44 | 3.94 | |
| | S.C. | -30 | -61 | 38 | 3.94 | |
| Blind | | | | | | |
| L middle occipital gyrus | 75 | -30 | -91 | -7 | 3.36 | <.015^ |
| Blind > Sighted | | | | | | |
| L middle occipital gyrus | 56 | -36 | -85 | -7 | 3.28 | .02^ |
| R middle occipital gyrus | 33 | 27 | -82 | 14 | 3.17 | .026^ |
| Seed in rPrecuneus | | | | | | |
| Blind | | | | | | |
| L middle frontal gyrus | 86 | -42 | 50 | 5 | 4.76 | .023* |
| L middle occipital gyrus | 234 | -24 | -76 | 26 | 4.55 | .001 |
| L superior parietal lobe | S.C. | -30 | -61 | 41 | 3.78 | |
| L superior parietal lobe | S.C. | -30 | -91 | 14 | 3.31 | |
| R middle occipital gyrus | 537 | 39 | -85 | 20 | 3.78 | <.001 |
| | S.C. | 33 | -88 | 2 | 3.69 | |
| R fusiform gyrus | S.C. | 24 | -52 | -16 | 3.75 | |
| Blind > Sighted | | | | | | |
| L middle occipital gyrus | 159 | -27 | -79 | 23 | 4.44 | .007 |
| | S.C. | -30 | -91 | 17 | 3.63 | |
| | S.C. | -42 | -82 | 20 | 3.20 | |
| R middle occipital gyrus | 170 | 39 | -85 | 20 | 3.95 | <.001 |
| | S.C. | 39 | -85 | 5 | 3.59 | |
| R superior occipital gyrus | S.C. | 24 | -88 | 23 | 3.45 | |

Significance corrections are reported at the cluster level, unless otherwise specified; cluster size threshold = 50. FWE, family-wise error; L, left; R, right; S.C., same cluster.

*Brain activations significant after FWE voxel-level correction over the whole brain. ^Brain activation significant after FWE voxel correction over a small spherical volume (10-mm radius) at peak coordinates for right/left MOG.

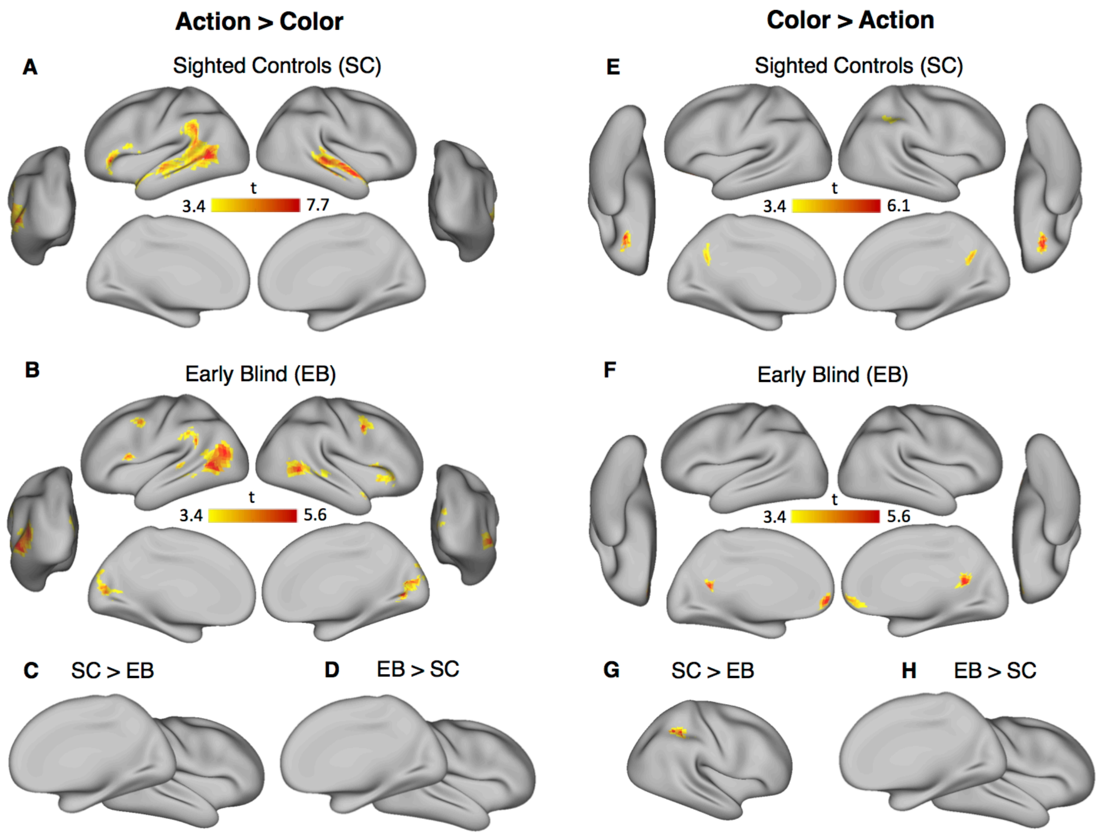


**Figure S1.** Regional BOLD responses are depicted over multiplanar slices and renders of the MNI-ICBM152 template. Suprathreshold cluster ($P<0.001$ uncorrected, for illustration only) showing brain activity for the contrasts (A) Action > Color in SC; (B) Action > Color in EB; (C) Action > Color & SC > EB, with no significant cluster; (D) Action > Color & EB > SC, with no significant cluster; (E) Color > Action in SC; (F) Color > Action in EB; (G) Color > Action & SC > EB; (H) Color > Action & EB > SC, with no significant cluster.

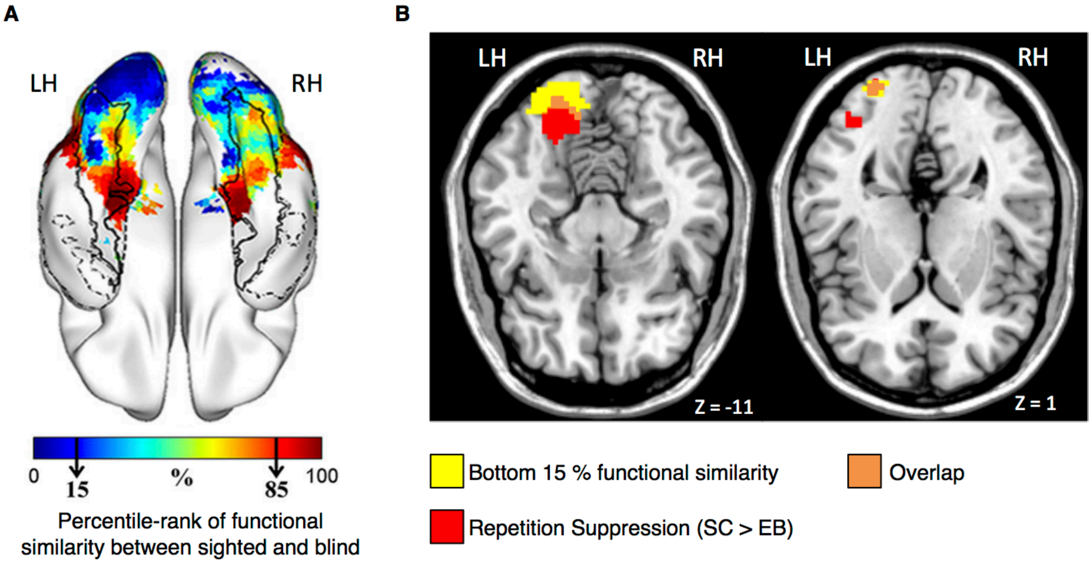


**Figure S2. Comparison with previous results.** (A) Percentile rank map of Pearson correlation coefficients of functional response patterns between blind and sighted groups (adapted from Wang et al. 2015). The figure shows that functional similarity of the VOTC between sighted and blind increases along a posterior-to-anterior gradient. Posterior occipital cortex has a different functional fingerprint in sighted and blind. Color bar represents the percentile rank value. Warmer colors represent greater between-group similarity. Black arrows in the color bar indicate the bottom and the top 15% similarity; (B) Areas showing the bottom 15% similarity between sighted and blind in Wang et al. experiment, and the areas showing greater repetition suppression in sighted than blind in our experiment ($p < .05$, FEW-corrected), and their overlap, are displayed on two axial views of the brain at z=-11 (Lingual Gyrus) and z=1 (Middle occipital Gyrus);

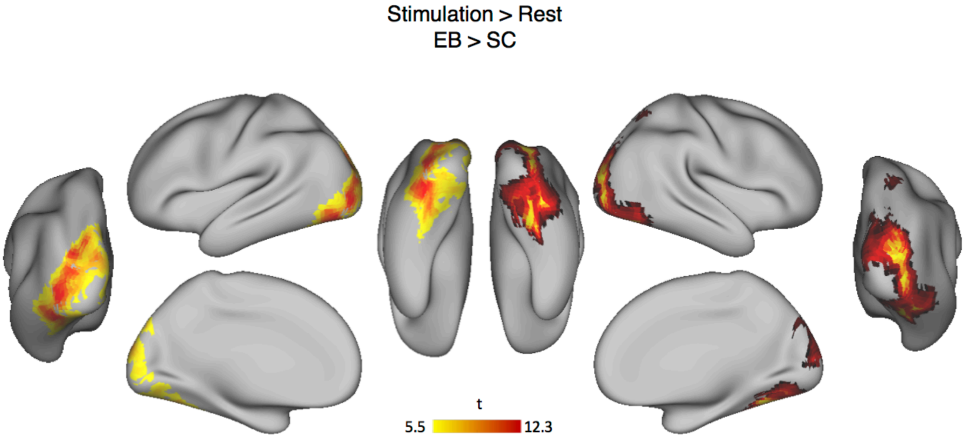


**Figure S3.** Whole brain results for the contrast Stimulation > Rest in blind more than sighted. FWE corrected at voxel level ($p < .05$). During the task (conceptual retrieval of actions and colors similarity) the occipital cortex of blind people is more active compared to the occipital cortex of sighted people. Although there is no evident category-specificity (e.g., preference for action or color concepts), our PPI analysis (see main manuscript) suggests that part of the occipital cortex in the blind works in concert with semantic hubs (pMTG, precuneus) during conceptual retrieval.

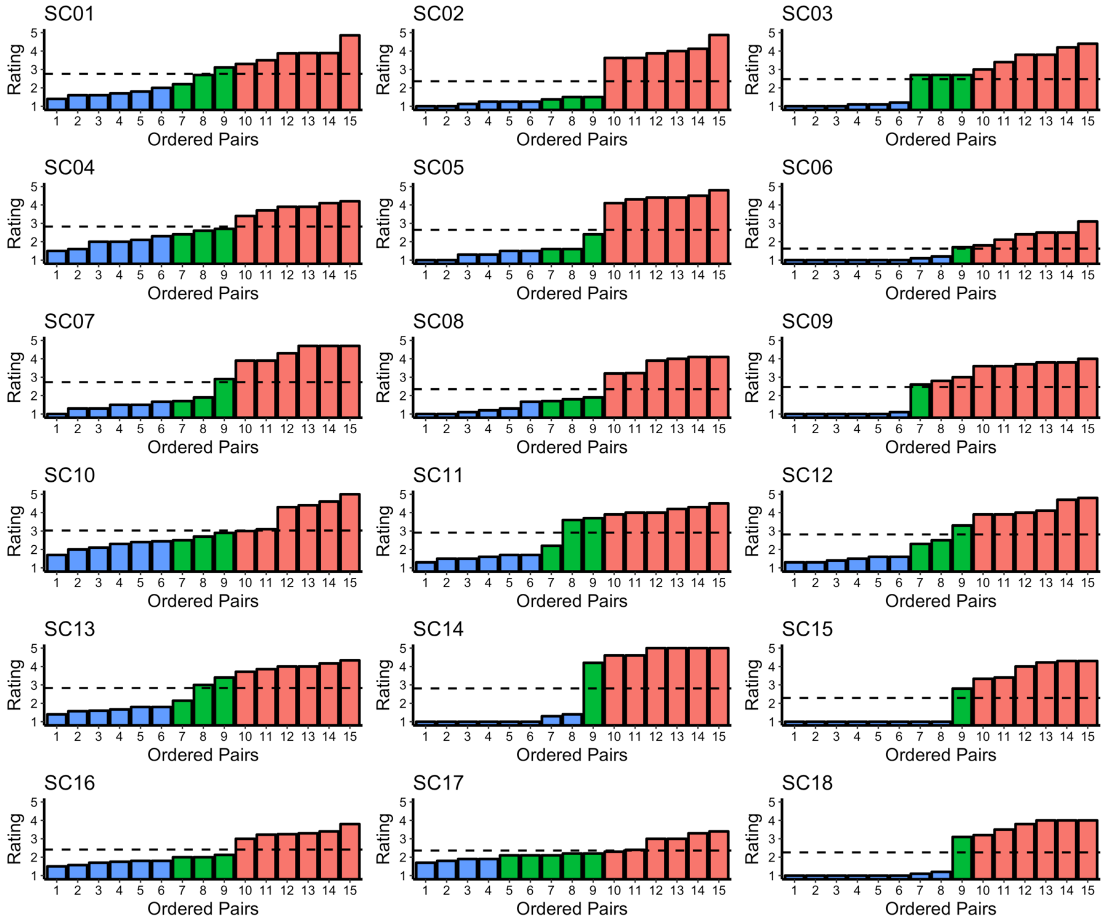


**Figure S4.** Action pairs divided by Similarity ratings in Sighted. In blue dissimilar, green medium and red similar pairs. Dashed line is the average rating for that given subject. Note that dissimilar pairs are always below average and similar pairs always above average.

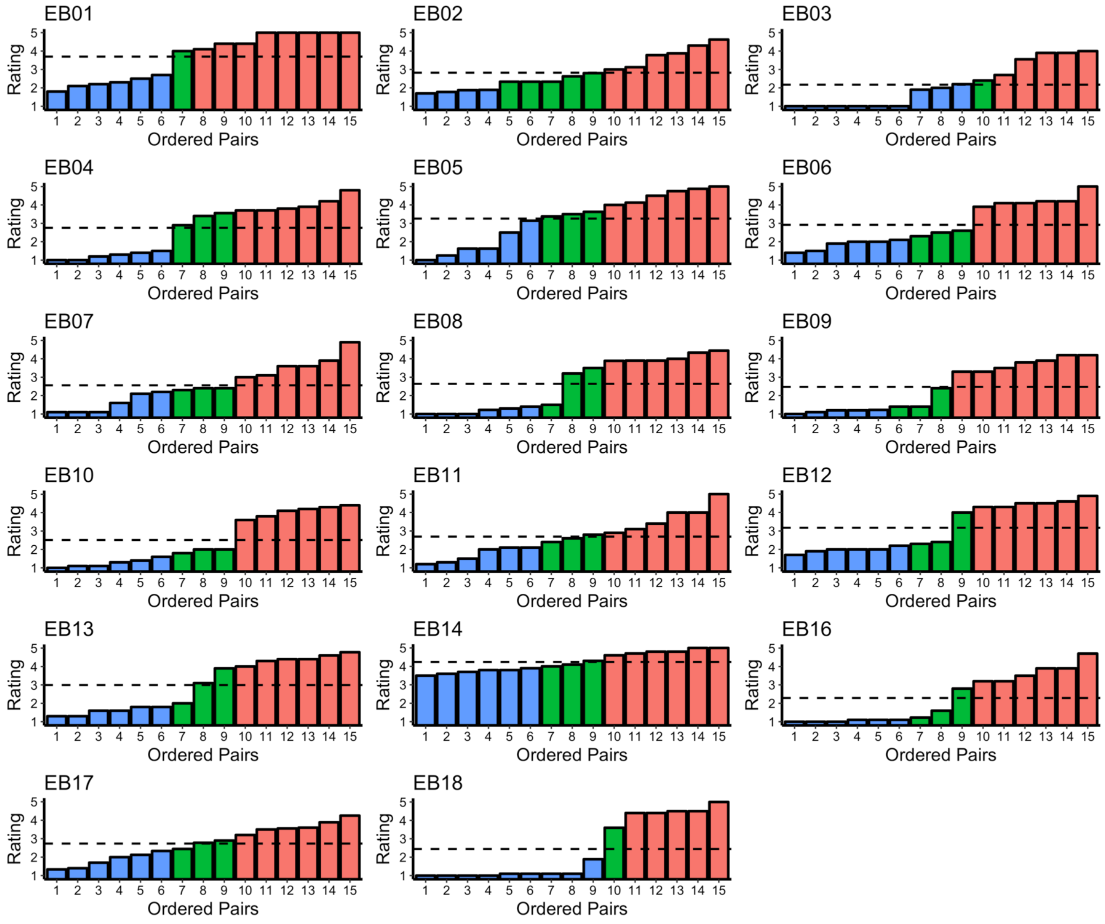


**Figure S5.** Action pairs divided by Similarity ratings in Blind. In blue dissimilar, green medium and red similar pairs. Dashed line is the average rating for that given subject. Note that dissimilar pairs are always below average and similar pairs always above average.

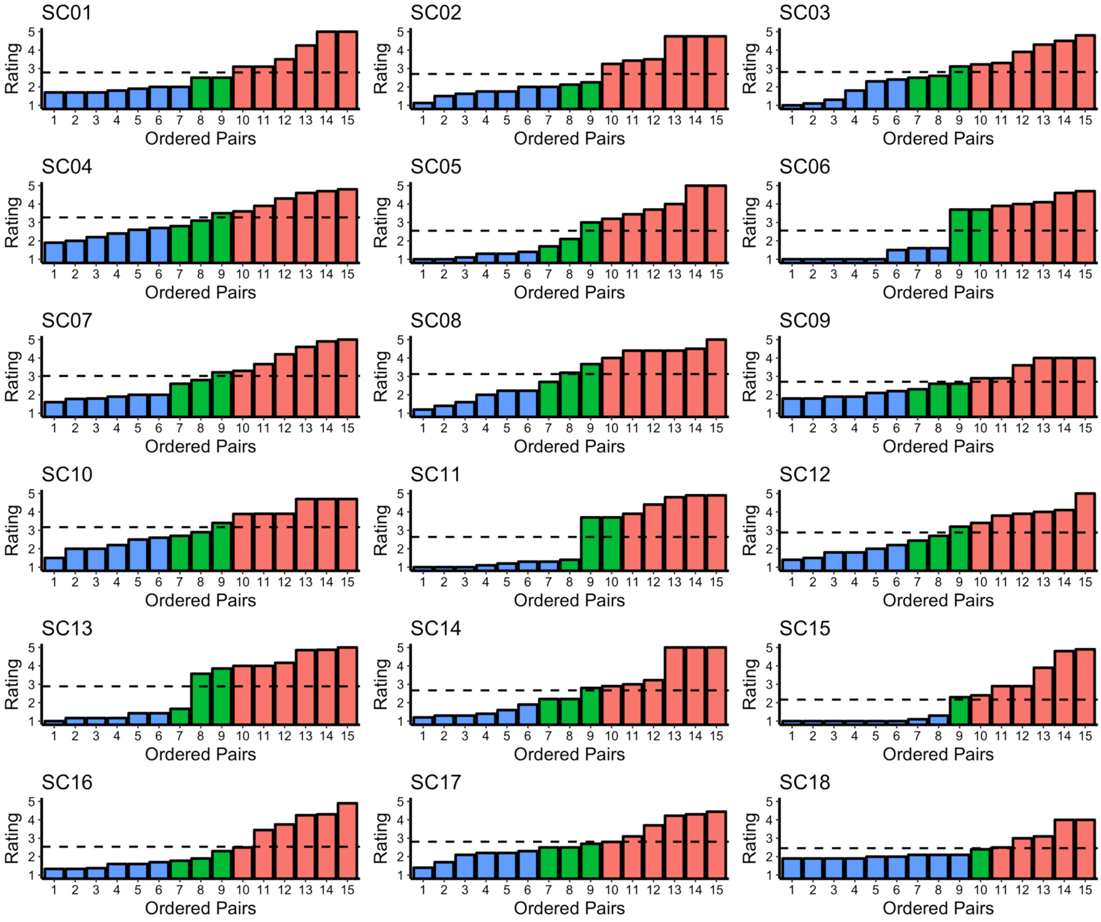


**Figure S6.** Color pairs divided by Similarity ratings in Sighted. In blue dissimilar, green medium and red similar pairs. Dashed line is the average rating for that given subject. Note that dissimilar pairs are always below average and similar pairs always above average.

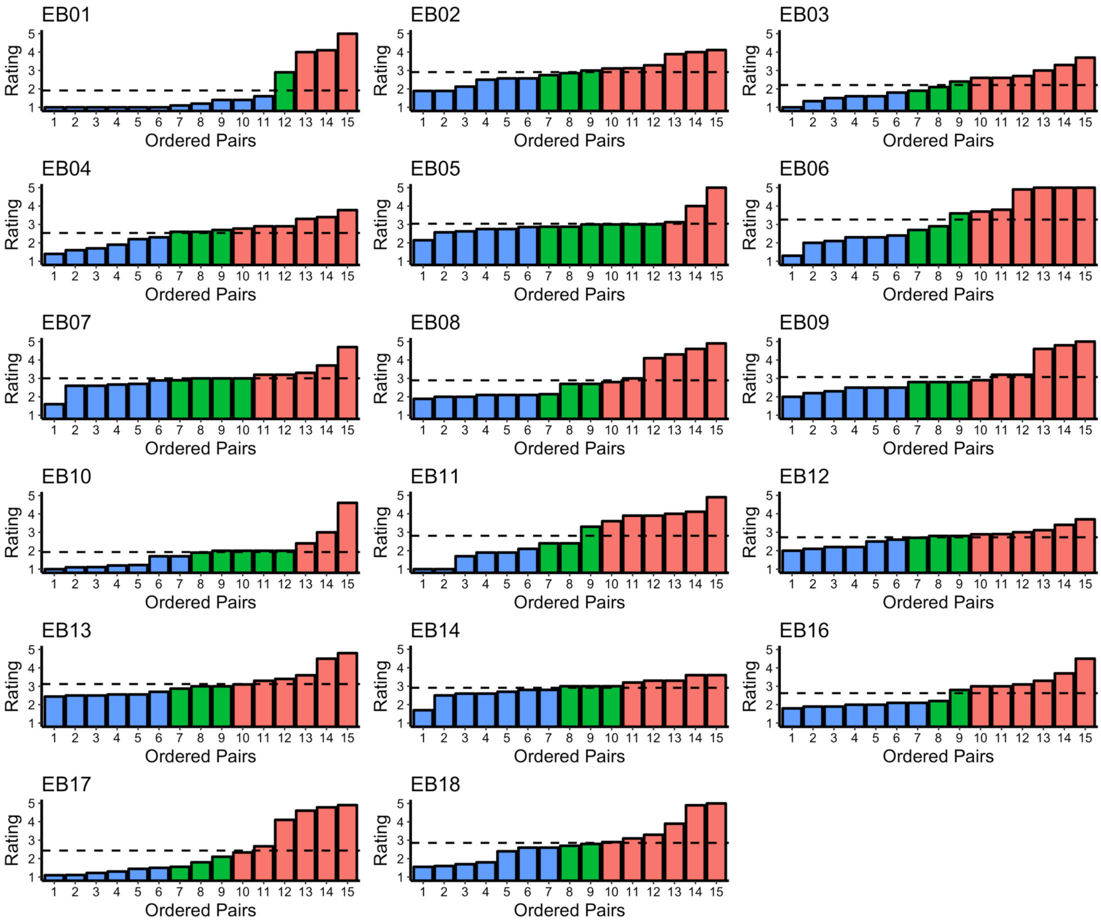


**Figure S7.** Color pairs divided by Similarity ratings in Blind. In blue dissimilar, green medium and red similar pairs. Dashed line is the average rating for that given subject. Note that dissimilar pairs are always below average and similar pairs always above average.

**Additional ROI analysis for PPI**

For the original PPI analysis, in each individual, time series of activity (principal eigenvariate) were extracted from a 8 mm sphere centered on the nearest local maxima to the identified peaks in the second-level analysis (seed region). However, centering the sphere on the peak itself does not change the ROI Analysis results:

PPI with seed in the lpMTG revealed an increase of action-selective functional connectivity in both occipital ROIs of blind people compared to their sighted counterpart (lMOG: $t= 3.33$, $p= 0.034$; rMOG: $t=3.21$, $p=0.043$). Similarly, PPI with seed in the precuneus revealed an increase of color-selective functional connectivity in the occipital cortices of blind compared to sighted participants (lMOG: $t= 3.44$, $p= 0.028$; rMOG: $t=3.81$, $p=0.012$).
